# eccDB: a comprehensive repository for eccDNA-mediated chromatin contacts in multi-species

**DOI:** 10.1101/2022.09.22.509011

**Authors:** Min Yang, Bo Qiu, Guo-You He, Jian-Yuan Zhou, Hao-Jie Yu, Yu-Ying Zhang, Yan-Shang Li, Tai-Song Li, Jin-Cheng Guo, Xue-Cang Li, Jian-Jun Xie

## Abstract

The role of extrachromosomal circular DNA (eccDNA) has been highlighted. More recently, eccDNA-chromosome interactions were identified, suggesting a potential role of eccDNA in transcriptional regulation. Several databases currently provide valuable resources for the study of eccDNAs. However, these databases are primarily focused on Human eccDNAs and do not provide analysis of eccDNA-chromosome interaction and eccDNA gene expression in different tissues. Herein, to further integrate available resources for eccDNA data across multiple species, we developed the eccDB database. The current version of eccDB has collected a total of 1,317,182 eccDNAs in 424 samples from four species (homo sapiens, mus musculus, saccharomyces cerevisiae, and arabidopsis thaliana). eccDB provides regulatory and epigenetic information on eccDNA, including typical enhancers, super-enhancers, transcription factors, DNA methylation positions, risk SNPs, expression quantitative trait locus, chromatin accessibility regions, and chromHMM states. In particular, eccDB provides eccDNAs intrachromosomal and interchromosomal interaction analysis to predict the transcriptional regulatory functions of eccDNA. Moreover, eccDB identifies eccDNAs from unknown DNA sequences and analyzes the functional and evolutionary relationships of an eccDNA among different species. Overall, eccDB offers web-based analytical tools and a comprehensive resource for biologists and clinicians to decipher the molecular regulatory mechanisms of eccDNA. eccDB is freely available at http://www.xiejjlab.bio/eccDB

## Introduction

Circular DNA is a form of DNA molecule that is widely found in nature. For example, bacterial plasmids or yeast mitochondrial DNA are circular DNA molecules. A special class of circular DNA molecules has been found in Eukaryotes. These circular DNA molecules separated or detached from the normal genome, free from the chromosomal genome. Since they are circular DNA molecules that exist outside the chromosome, they are called extrachromosomal circular DNA (eccDNA) (HOTTA and BASSEL 1965; Liao et al. 2020; Zuo et al. 2022).

Recent studies have shown that eccDNAs tend to be enriched during tumor and aging, eccDNAs are involved in tumorigenesis and development in a specific way. eccDNAs are widespread in most cancer cells and can carry complete genes, especially in tumor cells which often carry oncogenes (Paulsen et al. 2018). Oncogenes are highly enriched on amplified eccDNA, which promotes tumor genetic heterogeneity and accelerates tumor evolution (Kim et al. 2020). Early studies focused on the presence of eccDNA as an important form of oncogene amplification that may be important for the evolution of tumor cells, and suggested that its contribution to oncogene expression was mainly due to the increase in gene copy numbers (Kim et al. 2020; Kumar et al. 2020; Wang et al. 2021; Paulsen et al. 2018; Noer et al. 2022). Kohl *et al*. identified N-MYC gene amplification on eccDNA from neuroblastoma. Zhou YH *et al*. have found *EGFR* gene amplification on eccDNA in glioblastoma, and eccDNA-containing glioblastoma cells exhibited greater aggressiveness. In addition, approximately 30% of human epidermal growth factor receptor 2 (HER2)-positive breast cancers exhibit eccDNA HER2 amplification, and this eccDNA carrying the *HER2* gene is relatively conserved in drug-resistant tumors (Wang et al. 2021). Through computational analysis of whole genome sequencing data, Kim H *et al*. showed that eccDNAs amplification resulted in higher levels of oncogene transcription and higher expression of oncogenes on eccDNA. eccDNA-based oncogene amplification is common in cancer and leads to poor prognosis in patients with many cancer types (Kim et al. 2020).

Furthermore, eccDNAs have important transcriptional regulatory functions and show significantly enhanced chromatin accessibility (Wu et al. 2019). Zhu Y *et al*. used ChIA-PET and ChIA-Drop to elucidate the transcriptional interactions and regulation of eccDNAs in glioblastoma and prostate cancer cells. Through multi-genomic analysis of eccDNA-associated chromosomal genes, super-enhancer (SE) features, and RNA expression, these results confirm that eccDNA can act as a mobile transcriptional SE element to promote tumor progression. They applied ChIA-PET assays to pull down RNAPII-associated chromatin and identified eccDNA-chromosome interactions (Zhu et al. 2021). Overall, eccDNAs have important functions: eccDNAs carry genes that are highly expressed in cancer with poor prognosis and eccDNAs interact with chromosomes, and play potential roles in transcriptional regulation.

Several eccDNA databases have been established, such as CircleBase (Zhao et al. 2022) and eccDNAdb (Peng et al. 2022). These two databases provide valuable resources for the study of human eccDNAs. Both databases provide basic information about eccDNAs, such as chromosomal loci. CircleBase also provides eccDNAs intrachromosomal interactions, and eccDNAdb offers the analysis of eccDNAs gene expression and survival in human cancer. However, existing databases do not provide analysis of eccDNAs interchromosomal interactions and eccDNA genes expression in different tissues. Furthermore, existing databases do not provide relevant explorations of eccDNAs sequence similarity in multi-species.

To this end, we have developed a comprehensive repository for eccDNA-mediated chromatin contacts in multi-species (eccDB, http://www.xiejjlab.bio/eccDB). eccDB focuses on providing a large number of available resources for eccDNA across multiple species, including Homo sapiens, Mus musculus, Saccharomyces cerevisiae, and Arabidopsis thaliana. The current version of eccDB collects 767,981 eccDNAs from 396 human samples (hg38), 372,811 eccDNAs from 12 mouse samples (mm10), 2,081 eccDNAs from 11 Saccharomyces cerevisiae samples (sacCer3), and 174,309 eccDNAs from 5 arabidopsis thaliana samples (tair10). eccDB provides six query methods for searching eccDNAs. eccDNAs basic information, regulatory regions (typical enhancers (TEs), SEs, chromatin accessibility regions, chromHMM states), genetic and epigenetic information (transcription factors (TFs), risk SNPs, expression quantitative trait locus (eQTLs), DNA methylation positions) are available in eccDB. eccDNA genes transcriptional level, survival analysis, GO term functional enrichment analysis and KEGG pathway annotation are also provided. In addition, eccDB provides eccDNAs intrachromosomal and interchromosomal interaction analysis. This analysis can identify eccDNAs and genome-wide chromosomal interactions to discover eccDNA as mobile SEs for transcriptional activation of chromosomal genes. At last, eccDB can help users to analyze whether the sequences are similar to eccDNAs by BLAST + blastn (Camacho et al. 2009). Overall, eccDB is a comprehensive repository for storing, browsing, searching, and analyzing. Therefore, eccDB is a powerful platform to help users gain insight into the potential biological functions of eccDNAs.

## Results

### Global view of eccDB

eccDB is a comprehensive multi-species eccDNA database with multiple functions, such as browse, search, analysis, blast, statistics, and download. eccDB allows users to browse eccDNA sample information and provides multiple query methods. eccDB provides eccDNA basic information, eccDNAs gene expression analysis, survival analysis, GO term functional enrichment analysis, and KEGG pathway annotation. Besides, we provide sequence similarity analysis of eccDNA in four species. Notably, eccDB also provides eccDNA intrachromosomal and interchromosomal interaction analysis, identifying eccDNA and genome-wide chromosomal interactions which indicate potential transcriptional regulation information (Figure 1).

**Figure 1.**
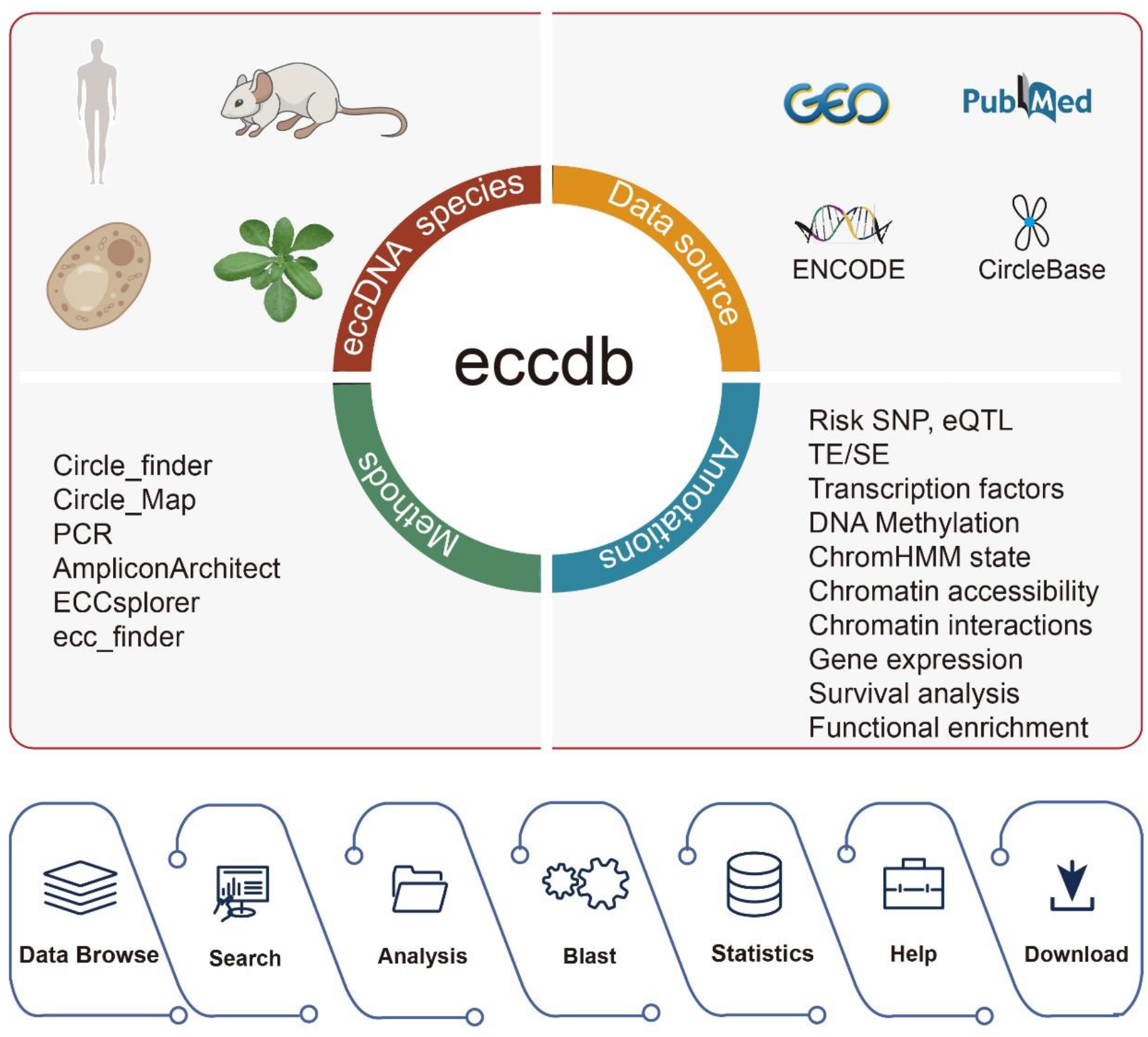
eccDB content and construction. eccDB collects eccDNA from Homo sapiens, Mus musculus, Saccharomyces cerevisiae, and Arabidopsis thaliana through a combination of eccDNA identification tools. eccDB annotates eccDNA regulatory elements, and genetic and epigenetic information including TE/SE, Chromatian accessibility, ChromHMM state, TFs, DNA methylation, Risk SNP, and eQTL. eccDB provides gene expression analysis, survival analysis, GO term functional enrichment analysis, and KEGG pathway annotation of eccDNAs. The eccDB provides sequence similarity analysis of eccDNA for four species and provides intrachromosomal and interchromosomal interaction analysis.

### Database usage and access

#### Data browse

eccDB provides a friendly browsing interface for users to browse sample information by species. Taking human eccDNA samples as an example, the left side of the “Browse” page provides “Data Source”, “Biosample Type”, “Disease Type” and “Tissue Type” to filter the information on the right side (Figure 2B, left). The right side shows the basic information of the sample in the form of a table, including “Sample_ID”, “Sample_type”, “Sample_name”, “Tissue_type”, “Disease_type”, “Method”, “EccDNA_number”, etc (Figure 2B, right). Users can click on “Sample_ID” to view specific information about the sample and the list of eccDNA identified in that sample (Figure 2E).

**Figure 2.**
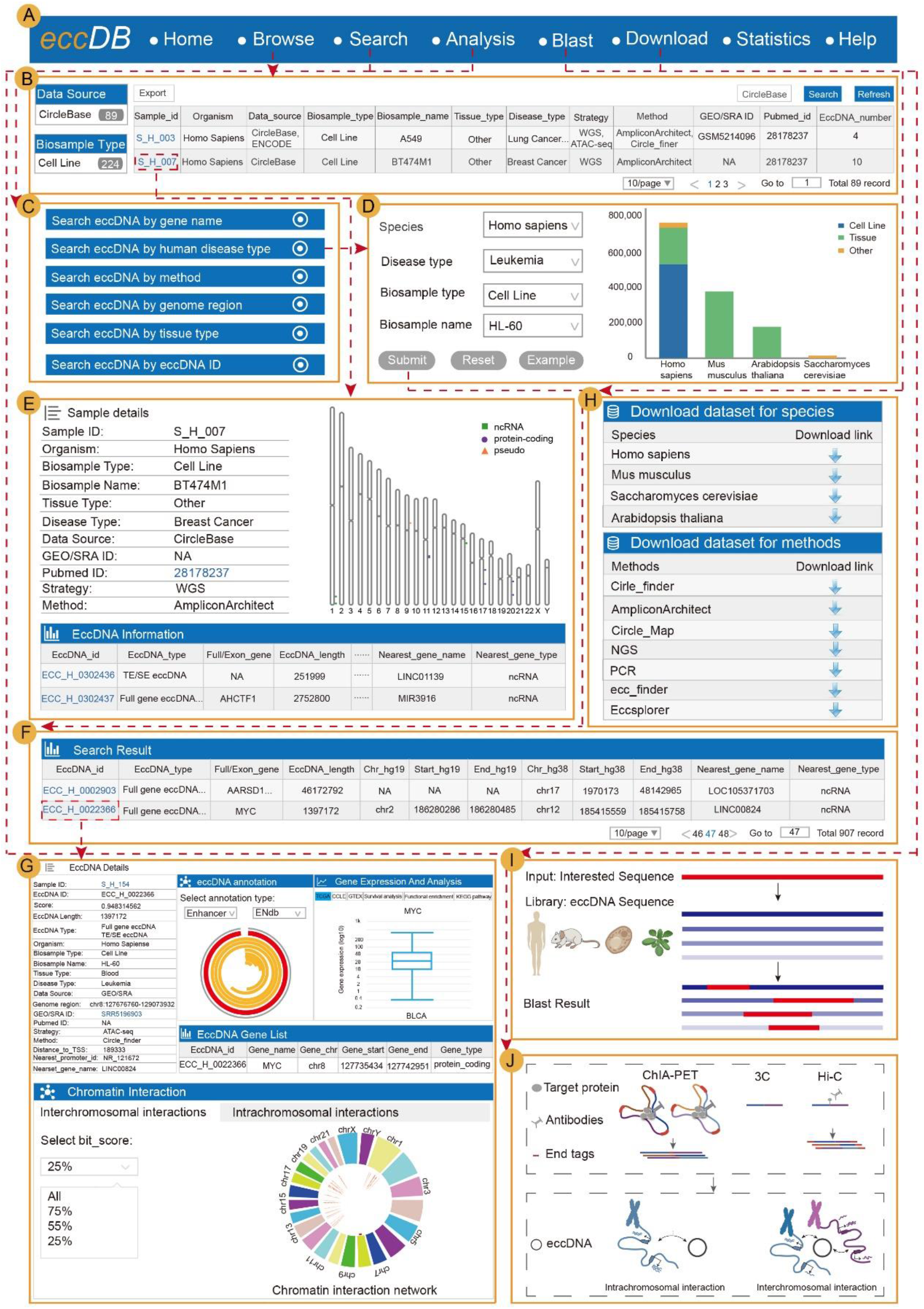
The main functions and usage of eccDB. (A) The navigation bar of eccDB. (B) The eccDB “Browse” page to view sample brief information in four species. (C) The eccDB provides six query methods for searching eccDNA: “Search eccDNA by gene name”, “Search eccDNA by human disease type”, “Search eccDNA by method”, “Search eccDNA by genome region”, “Search eccDNA by tissue type” and “Search eccDNA by eccDNA ID”. (D) Taking “Search eccDNA by human disease type” as an example, users select the disease type, biosample type, and biosample name of the human of interest. (E) This interface describes sample detail information and the list of eccDNA identified in this sample. (F) List of eccDNA searched by human disease type (Disease type: Leukemia; Biosample type: Cell Line; Biosample name: HL-60). (G) This interface is the eccDNA details page and describes the eccDNA sample information, eccDNA type, eccDNA gene list, eccDNA gene expression analysis and analysis (including survival analysis, GO term functional enrichment analysis, and KEGG pathway annotation), eccDNA chromatin interaction, and eccDNA annotation information (TEs, SEs, Chromatin accessibility regions, ChromHMM states, TFs, DNA methylation positions, Risk SNPs, eQTLs). (H) The “Download” allows users to download eccDNAs information by species, or eccDNAs information by methods. (I) The eccDB helps users analyze whether the nucleic acid sequence of interest has sequence similarity to eccDNA in four species. (J) The eccDB provides eccDNA intrachromosomal and interchromosomal interactions analysis.

#### A search interface for retrieving eccDNA

eccDB provides six query methods to search eccDNAs information, including “Search eccDNA by gene name”, “Search eccDNA by human disease type”, “Search eccDNA by method”, “Search eccDNA by genome region”, “Search eccDNA by tissue type”, “Search eccDNA by eccDNA ID” (Figure 2C). As an example, we searched for HL-60 cell lines in Leukemia by “Search eccDNA by human disease type” (Figure 2D), and a summary of the search results was displayed on the search results page (Figure 2F). The table described the eccDNAs statistics including EccDNA_ID, EccDNA_type, Full/Exon_gene, EccDNA_length, chromosome locus, Nearest_gene_name, Nearest_gene_type.

Users can click on “EccDNA_ID” to view detailed information about each eccDNA (Figure 2F). In addition to the brief information displayed on the search results page, users can also obtain the sample_type, sample_name, disease_type, tissue_type, and identification method for each eccDNA in eccDB (Figure 2G, top left corner). Additionally, eccDB also provides the sequence of eccDNAs. Specifically, users can select genes of interest in the “EccDNA Genes” module. eccDB will visualize the expression and survival analysis of eccDNA genes, GO term functional enrichment analysis, and KEGG pathway annotation (Figure 2G, upper right corner). Furthermore, eccDB visualizes genes through Integrative Genomics Viewer (IGV, https://igv.org/). eccDB also provides eccDNAs regulatory element information and epigenetic information, including TEs, SEs, etc. (Figure 2G, upper part of the ring diagram). Noteworthy is that eccDB shows the potential interchromosomal interaction and intrachromosomal interaction of eccDNA through tables and networks (Figure 2G, lower part).

#### eccDNA chromatin interaction analysis

In the “Analysis”, eccDB provides both eccDNAs intrachromosomal interaction analysis and interchromosomal interaction analysis. First of all, users enter a gene name or a chromosome region. Secondly, users select the species, tissue type, interaction type, and blast threshold to be analyzed, and click the “Analysis” button. Finally, eccDB will automatically analyze the eccDNA with potential interactions with the user-entered genes.

Taking a gene as an example, the analysis principle is as follows: Firstly, eccDB finds the chromosomal region (Interactor_chr, Interactor_start, Interactor_end) of the gene entered by the user. Then matches to the chromosomal fragment (Subject_region) provided by 4DGenome (https://4dgenome.research.chop.edu/) (Teng et al. 2015) that interacts with the “Interactor_region”. Secondly, using BLAST + blastn (Camacho et al. 2009) to find out the “EccDNA_region” with a similar sequence to the “Subject_region”. Thirdly, matching the corresponding to “EccDNA_ID” based on “EccDNA_region”. Finally, eccDB infers that the user-entered gene has potential interactions with “EccDNA_region”.

#### Blast analysis

In the “Blast”, eccDB provides sequence similarity analysis using BLAST + blastn (Camacho et al. 2009). eccDB constructs eccDNA nucleic acid sequences library in four species (human, mouse, saccharomyces cerevisiae and arabidopsis thaliana). After the user enters a sequence or uploads a “. fa” file (less than 5MB, the sequence or file should be a nucleic acid sequence of interest to the user), eccDB will randomly assign a “Job ID”. And then, eccDB applies the BLAST + blastn to compare the entered sequence with the nucleic acid library according to the parameters set by the user. Finally, the user can input the “Job ID” to obtain the analysis result. Users can analyze whether interest nucleic acid sequences have sequence similarity with eccDNA in four species. In general, eccDB is beneficial to help users explore homologous sequences in different species (Figure 2I).

#### Data download and statistics

On the “Download”, users can download eccDNAs information by species, or eccDNAs information by method (Figure 2H). The “Statistics” provides statistical charts for five panels and a statistical table. The five statistical charts include the number of eccDNA identified per sample, the number of eccDNA identified by each method, the number of eccDNA identified per data source, the number of eccDNA on chromosomes, and the number of eccDNA identified for each species. And the statistical table shows the differences between eccDB and other eccDNA databases in terms of the number of samples, the number of eccDNA, and database functions, as shown in Supplementary Table S1.

#### Case studies

In the following two case studies, we illustrate how eccDB can be used to select candidate eccDNA in different research for further experiments.

#### Case study for eccDNA chromosomal interaction

Multi-genomic analysis of eccDNA-associated chromosomal genes, SE features, and RNA expression demonstrated that eccDNAs can act as mobile transcriptional SE elements to promote tumor progression and showed a potential transcriptional regulatory mechanism for synthetic aneuploidy. Zhu Y *et al*. adopted ChIA-Drop assays and identified eccDNA-chromosome interactions in the PC3 human prostate cancer cell line (Zhu et al. 2021).

To exemplify the accuracy of eccDB for collecting information on eccDNA-chromosome interactions, we used the “Search eccDNA by human disease type” (Disease type: Prostate cancer; Biosample type: Cell Line; Biosample name: PC3) as search criteria (Figure 3B). The output table first shows brief annotation information of all eccDNAs identified in PC3 (Figure 3C). Taking “ECC_H_0302418” as an example, we can see that the eccDNA “ECC_H_0302418” carries the *MYC* gene. SEs are a large cluster of transcriptionally active enhancers that play a key role in the identity of human cells in health and disease, driving gene expression (Hnisz et al. 2013). eccDB shows that “ECC_H_0302418” has 227 SE regions from SEdb database (Figure 3E), 49 chromatin accessibility regions, and 231 transcription factor regions, which has potentially high transcriptional activity. Similarity analysis of DNA sequence is a necessary process to compare unknown DNA sequences with known ones for inferring the functions of unknown ones (Wang et al. 2010). Therefore, eccDB provides chromosomal interaction prediction based on DNA sequence similarity and chromosome interaction data (including ChIA-PET, 3C, and Hi-C). And the eccDNA “ECC_H_0302418” has 238 interchromosomal interactions and 7 intrachromosomal interactions.

**Figure 3.**
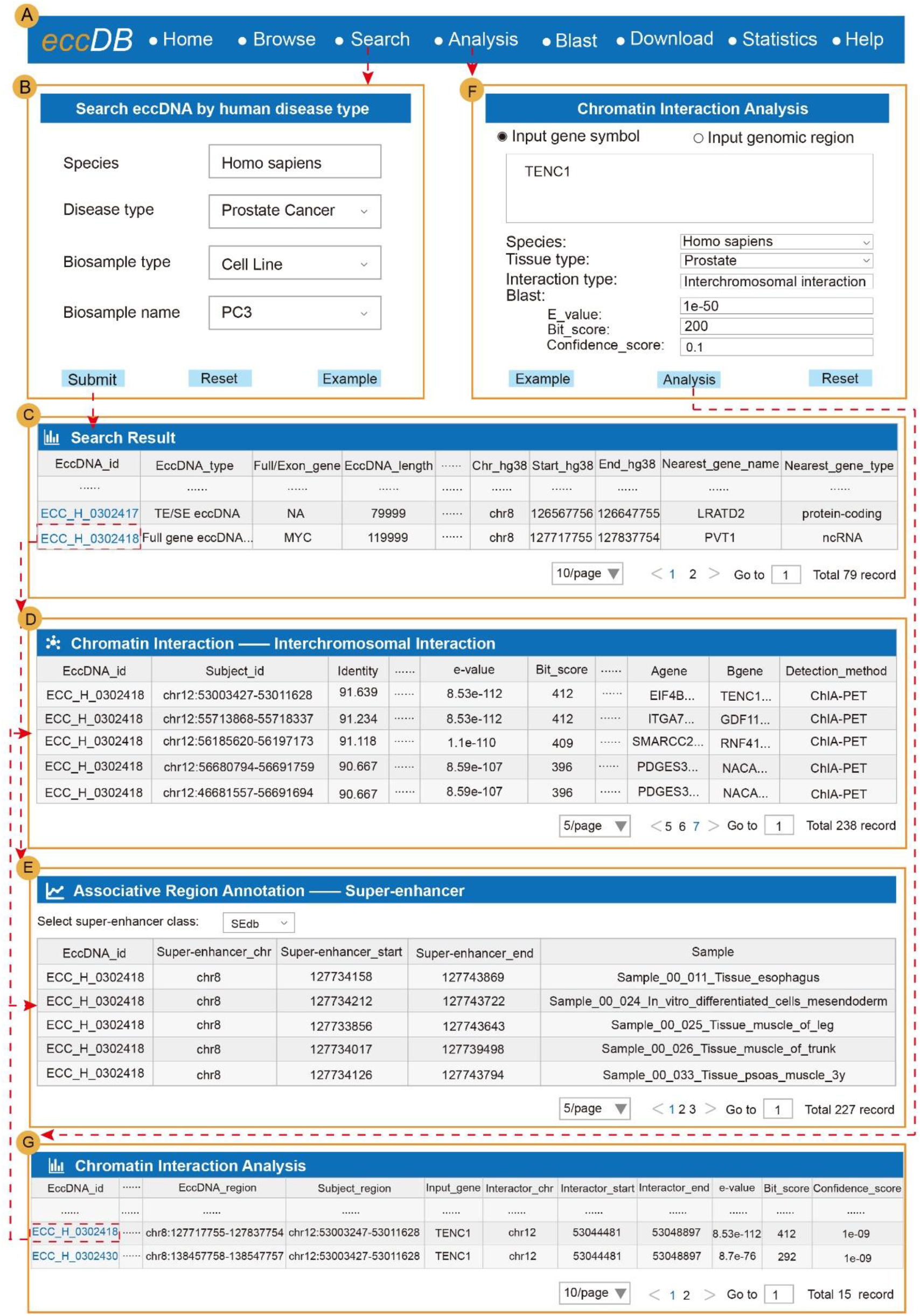
Case study for eccDNA chromosomal interaction. (A) The eccDB navigation bar. (B) Search eccDNA by human disease type (Disease type: Prostate Cancer; Biosample type: Cell Line; Biosample name: PC3). (C) List of eccDNA searched by human disease type (set search criteria by Figure 3B). (D) List of “ECC_H_0302418” interchromosomal interaction. (E) List of “ECC_H_0302418” super-enhancer from SEdb. (F) The chromosome interaction analysis of *TENC1* gene, set species as human, tissue type as prostate, interaction type as interchromosomal interaction, and Blast as default value. (G) List of “TENC1” interchromosomal interaction.

Additionally, eccDB identified sequence similarity between “ECC_H_0302418” and the *EIF4B* gene (chr12: 53003427-53011628, e_value = 8.53e-112, bit_score = 412). In “Chromatin Interaction”, the table of interchromosomal interaction shows that the *TENC1* and *EIF4B* interaction was verified by ChIA-PET. Therefore, we inferred that “ECC_H_0302418” and *TENC1* have a potential interaction relationship based on sequence similarity (Figure 3D). The “ECC_H_0302418” also interacts with other genes, such as *RNF41, CSDE1, ATP5B*, and *CIRBP*. These genes can be queried in the list of genes interacting with eccDNA detected by Zhu Y *et al*. (Zhu et al. 2021).

Furthermore, when we sequentially entered genes *TENC1, RNF41, CSDE1, ATP5B*, and *CIRBP* on the “Analysis” page, set species as Homo sapiens, tissue type as Prostate, interaction type as interchromosomal interaction, and blast section as default parameters for analysis (Figure 3F). The result showed that all these genes interacted with “ECC_H_0302418”. This indicates that the results of the “Analysis” are mutually validated. And the “Analysis” can be used to predict the interaction between eccDNAs and chromosomes (Figure 3G).

#### Case study for eccDNA transcription

In cancer, oncogenes were usually amplified on eccDNAs, and pan-cancer analysis revealed that oncogenes encoded by eccDNAs were highly expressed in the tumor transcriptome (Paulsen et al. 2018; Kim et al. 2020). It is also reported that eccDNAs showed significantly enhanced chromatin accessibility (Wu et al. 2019).

In “Search eccDNA by human disease type”, we input Disease type: Glioblastoma; Biosample type: Cell Line; Biosample name: GBM39 (Figure 4B). Taking “ECC_H_ 0458354” as an example, we found that this eccDNA carried *LANCL2* gene, *VOPP1* gene and *EGFR* gene, of which *EGFR* is an oncogene (Figure 4C). eccDB showed the expression of *EGFR* (TCGA) in Glioblastoma (GBM) tumor samples with box plots (Figure 4D, left). Furthermore, eccDB provided survival analysis showing that glioblastoma patients with *EGFR* high-expression have a lower survival rate (Figure 4D, right). Since the “*EGFR*” gene is on “ECC_H_0458354”, we infer that cancer patients carrying “ECC_H_0458354” may have a poorer prognosis.

**Figure 4.**
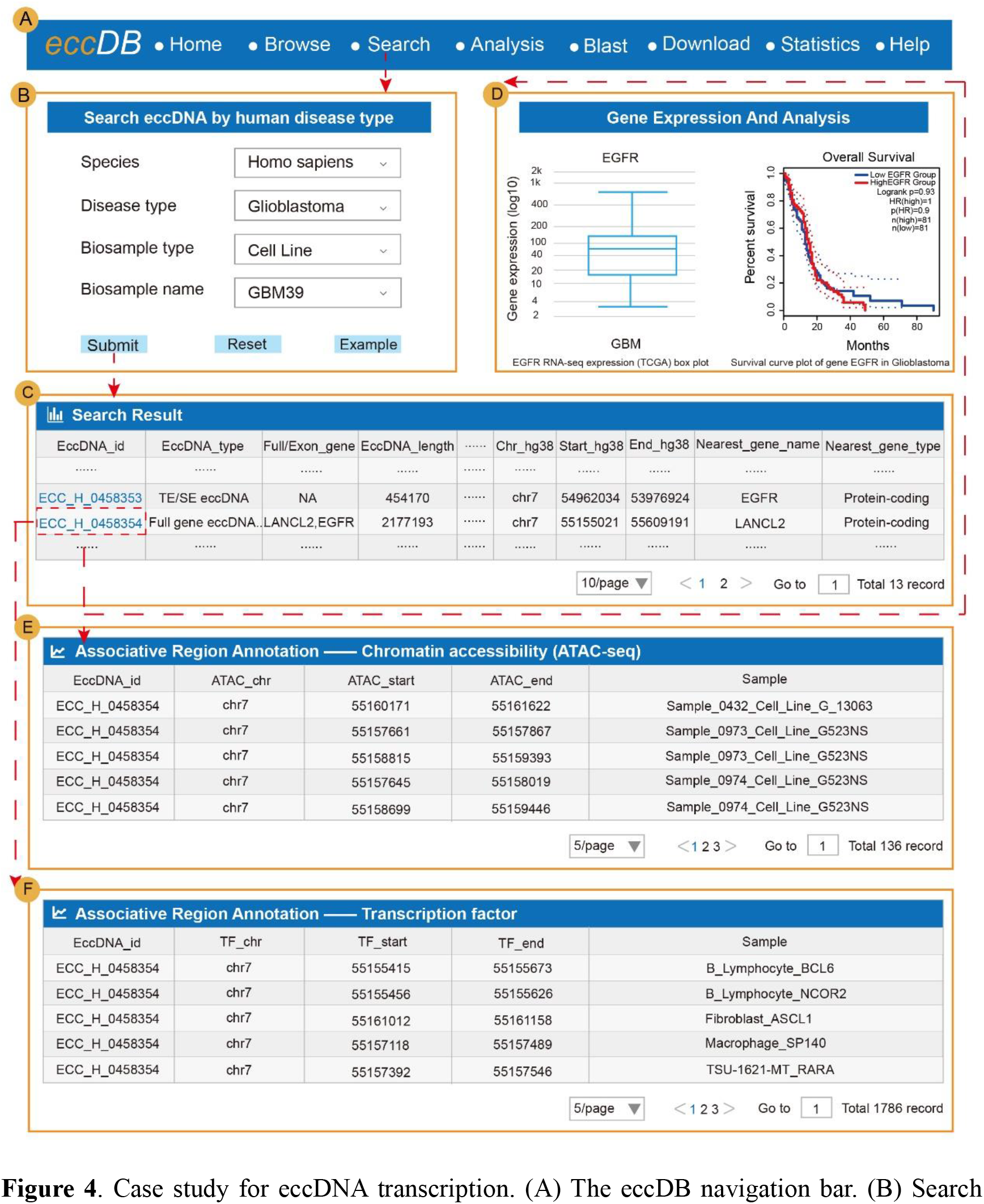
Case study for eccDNA transcription. (A) The eccDB navigation bar. (B) Search eccDNA by human disease type (Disease type: Glioblastoma; Biosample type: Cell Line; Biosample name: GBM39). (C) List of eccDNA searched by human disease type (set search criteria by Figure 4B). (D) On the left, the expression of *EGFR* (TCGA) in Glioblastoma (GBM) tumor samples with box plots, and the survival curve of *EGFR* in GBM on the right. (E) Information about the chromatin accessibility regions of “ECC_H_0458354”. (F) Information about the transcription factors of “ECC_H_0458354”.

Studies have shown that *EGFR* gene is highly expressed in GBM and amplification of the *EGFR* gene was associated with a poorer prognosis (Layfield et al. 2006; Xu and Li 2018). Interestingly, Wu S *et al*. found that oncogenes encoded on eccDNA (including *EGFR, MYC, CDK4*, and *MDM2*) were highly expressed in the cancer genome (Wu et al. 2019). eccDNAs carried *EGFR* gene was identified in patient-derived GBM cell lines by Kumar P *et al*. The *EGFR* gene is amplified through the formation of eccDNA in GBM (Kumar et al. 2020). These studies indicate that eccDNAs contribute to the upregulation of *EGFR* which is also found in our database.

In addition, 136 chromatin accessibility regions and 1,786 transcription factors binding sites were found on “ECC_H_0458354” based on ATAC-seq data (Figure 4E) (Figure 4F), validating that this eccDNA shows significantly enhanced chromatin accessibility in cancer.

In summary, we provided two effective examples to illustrate how to use eccDB to identify potential chromosome interaction and predict eccDNA gene’s function. More importantly, these results further demonstrate the usefulness and value of eccDB.

## Discussion

It has been reported that eccDNA is closely associated with cancer, drug resistance, and aging, while eccDNA plays an important role in gene compensation and the tumor-specific profile of eccDNA is even more important for disease recognition and prognosis (Zuo et al. 2022). Besides, studies have shown that eccDNA occurs in clusters in cancer or acts as a mobile SE element in transcriptional regulatory (Zhu et al. 2021; Hung et al. 2021). More importantly, eccDNA has universal eccDNA-chromosome interactions (Zhu et al. 2021). With the development of eccDNA identification methods (e.g., Circle_finder (Paulsen et al. 2018), Circle_Map (Prada-Luengo et al. 2019), ecc_finder (Zhang et al. 2021), AmpliconArchitect (Deshpande et al. 2019), ECCsplorer (Mann et al. 2022)). And the in-depth study of eccDNA biological functions by biologists, there is an urgent need for an integrated database to consolidate basic eccDNA information (e.g., chromosomal loci, sample types, disease types.) and to mine the potential functions of eccDNA. Current eccDNA databases include CircleBase (Zhao et al. 2022) and eccDNAdb (Peng et al. 2022). These databases provide a valuable resource for eccDNA collections, providing partial functional annotation and visualization. However, the existing databases do not provide eccDNA-chromosomal interactions analysis, eccDNAs classification, and eccDNA gene function annotation.

Therefore, we developed eccDB, a comprehensive repository for eccDNA-mediated chromatin contacts in multi-species. Specifically, eccDB not only collects basal information on eccDNAs of four species (Homo sapiens, Mus musculus, Saccharomyces cerevisiae, and Arabidopsis thaliana), but also annotates eccDNAs regulatory elements (TEs, SEs, chromatin accessibility regions, chromHMM states), genetic and epigenetic information (TFs, risk SNPs, eQTLs, DNA methylation positions). In particular, eccDB classifies eccDNAs into four categories and annotates them with full gene, exon gene, and nearest gene, and provides eccDNA gene expression visualization, survival analysis in cancer samples, GO term functional enrichment analysis and KEGG pathway annotation. In addition, eccDB provides eccDNA sequence similarity analysis which furtherly help provide extensive chromatin interaction data (e.g., ChIA-PET, 3C, and Hi-C) for analyzing eccDNA interchromosomal and intrachromosomal interaction.

eccDB supports a user-friendly interface to query, analyze, browse and visualize detailed information about eccDNA. The advantages of eccDB include: (I) a comprehensive repository for eccDNA, (II) six query methods for eccDNA study, (III) eccDNA genes information, (IV) eccDB sequences similarity analysis, and (V) eccDNA interchromosomal interaction and intrachromosomal interaction analysis.

Although eccDB already provides a large number of multi-species eccDNAs information, we believe that as eccDNAs are furtherly studied, the dataset of eccDNAs will rapidly accumulate and their potential functions will be better understood. And we will continue to collect available resources and process this data to integrate a more comprehensive functional annotation of eccDNA. Existing eccDNA chromosome interaction recognition methods are scarce and simple, with low reproducibility. To improve the accuracy and convenience of eccDNA chromosome interaction recognition, we will consider combining machine learning to mine an eccDNA chromosome interaction recognition method. In future releases, we will provide more comprehensive information on eccDNA. Overall, eccDB can help users to gain insight into the mechanisms of occurrence and potential biological functions of eccDNA.

## Methods and materials

### eccDNA collection and identification

There are already multiple ways to identify eccDNAs, so we collected a large number of multi-omics data and six eccDNA identification methods (Circle_ finder, Circle_ Map, PCR, AmpliconArchitect, ECCsplorer, ecc_ finder). To collect human eccDNAs, we downloaded from a large number of Assay for Transposase-Accessible Chromatin with high-throughput sequencing (ATAC-seq) data (Kumar et al. 2020) and Circle-Seq data (Møller 2020) from GEO/SRA (Barrett et al.2013) and ENCODE (ENCODE Project Consortium. 2012) for the identification of eccDNA. A part of human eccDNAs were downloaded from CircleBase (Zhao et al. 2022). To collect mouse eccDNAs, we loaded extensive ATAC-seq data from GEO/SRA, then conducted an extensive literature search in PubMed (https://www.ncbi.nlm.nih.gov/pubmed/) and integrated the collection of mouse eccDNAs by literature mining. For saccharomyces cerevisiae and arabidopsis thaliana, we collected publicly available eccDNAs information by collection of literatures (Figure 1).

For ATAC-seq data, eccDNAs identification was performed using Cirlce_finder (https://github.com/pk7zuva/Circle_finder), which is a method to identify circular DNA from pair-end high-throughput sequencing data (Kumar et al. 2017; Dillon et al. 2015). First, we use circle_finder-pipeline-bwa-mem-samblaster.sh to map human ATAC-seq data to the human genome hg38 and mouse ATAC-seq data to the mouse genome mm10. Finally, length of eccDNA <50MB was retained.

For the Circle-Seq data, eccDNAs identification was performed using Circle-Map (https://github.com/iprada/Circle-Map), the algorithm accurately detects circular DNA formed from mappable and non-mappable regions of a genome (Prada-Luengo et al. 2019). First, we used BWA (Jung and Han 2022) to align the reads to the human genome hg38 downloaded from GENCODE (https://www.gencodegenes.org/) (Harrow J et al. 2012). Then SAMtools (Li et al. 2009) was used to index the results in SAM/BAM format to compare the results. Thirdly, Circle-Map Realign (Prada-Luengo et al. 2019) was used to identify the eccDNAs. Finally, the eccDNAs with “Split reads=0” in the result are deleted and the eccDNAs with length <50MB is retained.

In addition, eccDB also includes eccDNAs identified by other methods (including ecc_finder (Zhang et al. 2021), AmpliconArchitect (Deshpande et al. 2019) and ECCsplorer (Mann et al. 2022), etc.) from literatures (Zhang et al. 2021; Mann et al. 2022; Dillon et al. 2015; Shibata et al. 2012; Wang et al. 2021; Møller et al. 2015). Detailed sample numbers and eccDNA numbers for each species can be viewed in Table 1.

**Table 1.** The statistics about the sample type and eccDNA counts of eccDB

|  | Homo<br>sapiens | Mus<br>musculus | Saccharomyce<br>s cerevisiae | Arabidopsis<br>thaliana |
| --- | --- | --- | --- | --- |
| Sample types counts | 396 | 12 | 11 | 5 |
| eccDNA counts | 767,981 | 372,811 | 2,081 | 174,309 |
| TEs | 6,370,152 | 47,198 | 0 | 0 |
| SEs | 393,782 | 828 | 0 | 0 |
| chromatin accessibility regions | 39,586,081 | 0 | 0 | 0 |
| ChromHMM states | 37,288,007 | 0 | 0 | 0 |
| TFs | 28,500,150 | 6,294,670 | 739 | 0 |
| Risk SNPs | 1,515,093 | 0 | 0 | 0 |
| eQTLs | 10,115,157 | 0 | 0 | 0 |
| Methylation positions | 776,369 | 0 | 0 | 0 |
| Intrachromosomal interaction | 3,163,288 | 79,553 | 55,579 | 0 |
| Interchromosomal interaction | 29,454,494 | 1,417,698 | 204,030 | 0 |

### eccDNA genes annotation

eccDNAs are widely present in most cancer cells and carry complete genes, especially oncogenic genes. We classified eccDNA genes into “Full gene”, “Exon gene” and “Nearest gene”. The “Full gene” is a protein-coding gene which all exons in the eccDNA region, and the “Exon gene” is a protein-coding gene with some exons in the eccDNA region. The “Nearest gene” is a gene that is close to eccDNA. To obtain the “Full gene” and the “Exon gene”, reference genomes of four species were downloaded. The human genome hg38 and mouse genome mm10 from GENCODE (https://www.gencodegenes.org/; Harrow et al. 2012), the saccharomyces cerevisiae genome sacCer3 from UCSC (https://genome.ucsc.edu/), and the arabidopsis thaliana genome tair10 from ENSEMBL PLANTS (https://plants.ensembl.org/index.html). Then screened the protein-coding genes of each species. Afterward, the “Full gene” and the “Exon gene” of the four species were annotated using BEDTools intersect (v2.25.0) (Quinlan et al. 2010). Finally, we annotated the eccDNAs loci by HOMER (Heinz et al. 2010) to obtain the “Nearest gene” for eccDNAs.

### eccDNA classification

eccDNAs structural diversity contributes to the functional diversity of eccDNAs, we thereby divided eccDNAs into four categories: Full gene eccDNA, Exon eccDNA, TE/SE eccDNA, and Intergenic eccDNA (Ling et al. 2021; Møller et al. 2020). The “Full gene eccDNA” is an eccDNA that contains one or more complete protein-coding genes. The “Exon eccDNA” is an eccDNA with part exons of one or more protein-coding genes. The “TE/SE eccDNA” is an eccDNA with TE or SE within the eccDNA region. The “Intergenic eccDNA” is an eccDNA that is derived from intergenic regions.

### eccDNA genes expression and analysis

The presence of amplification of oncogenes and drug resistance genes on eccDNAs has been reported, and the high expression level of eccDNA genes can be inferred (Zuo et al. 2022). We downloaded human RNA-seq expression data form TCGA (Corces et al. 2018) in UCSC Xena (https://xenabrowser.net/datapages/), and GTEX (Carithers and Moore 2015) in the human protein atlas(https://www.proteinatlas.org/), and human RNA-seq expression data from CCLE (https://sites.broadinstitute.org/ccle/; Barretina et al. 2012), then visualized eccDNAs gene expression using box plots. In addition, survival analysis of eccDNAs gene expression in cancer samples was performed by GEPIA2 (Tang et al. 2019) in “Gene Expression and Analysis”. eccDB also provided GO term functional enrichment analysis and KEGG pathway annotation of the eccDNAs genes. The GO term functional enrichment analysis can be performed using “GO_Molecular_Function_2021”, #x201C;GO_Cellular_Component_2021”, and “GO_Biological_Process_2021” of R package enrichR. And the KEGG pathway annotation was performed using “KEGG_2021_Human” of R package enrichR (Figure 2G; Kuleshov et al. 2016).

### eccDNA nucleic acid sequence library building

Similar sequences usually have the same functions, eccDB provided the “Blast” function to discover whether the interest sequences file is similar to the eccDNAs sequences in different species. We used BLAST + Blastn to analyze sequence similarity. BLAST + is a tool to search for sequence similarity (Camacho et al. 2009). eccDB will post the eccDNAs chromosomal loci of human, mouse, saccharomyces cerevisiae, and arabidopsis thaliana to genome hg38, mm10, sacCer3, tair10 respectively through BEDTools getfasta (v2.25.0) (Quinlan et al. 2010). The human genome hg38 and mouse genome mm10 from GENCODE (https://www.gencodegenes.org/; Harrow et al. 2012), the saccharomyces cerevisiae genome sacCer3 from UCSC (https://genome.ucsc.edu/), and the arabidopsis thaliana genome tair10 from ENSEMBL PLANTS (https://plants.ensembl.org/index.html). Then BLAST + makeblastdb command was applied to build the eccDNA sequence library.

### eccDNA annotation information

To analyze the potential biological functions of eccDNA, eccDB provided regulatory elements, genetic and epigenetic annotations for eccDNAs, including TEs, SEs, TFs, DNA methylation positions, risk SNPs, eQTLs, chromatin accessibility regions, chromHMM states. For format and version consistency, liftOver (Karolchik et al. 2003) was used to convert genome versions for TEs, SEs, TFs, chromatin accessibility regions, and chromHMM states. DNA methylation positions, risk SNPs and eQTLs were used NCBI remap (http://www.ncbi.nlm.nih.gov/genome/tools/remap) to transform. The annotation of regulatory elements, genetic and epigenetic information of eccDNA is then done by BEDTools intersect (v2.25.0) (Quinlan et al. 2010). And the parameter is: bedtools intersect -a eccDNA.txt -b dataset.txt -F 1 -wa -wb > result.txt.

A total of 396,673 SEs from dbSUPER (Khan and Zhang 2016) and SEdb (Jiang et al. 2019) were downloaded, of which 65,884 SEs from 99 cells/tissues were obtained from dbSUPER, leaving 330,789 SEs from the 542 cells/tissues mentioned in SEdb. We downloaded 729 TEs from ENdb (Bai et al. 2020) and 6,456,640 TEs from SEdb. The chromatin accessibility regions data were all obtained from ATACdb (Wang et al. 2021), and these regions encompassed 1,493 samples of ATAC-seq data covering a wide range of cell/tissue types. We downloaded the TF ChIP-seq data from GREAP (Yang et al. 2022), whose TF data was provided by Cistrome (Mei et al. 2017) and Remap (Chèneby et al. 2018). The TF binding region information covers over 9,000 samples, including 57 tissue types and 3,328 TFs.

ChromHMM (Ernst and Kellis 2012) is an automated computational system for learning chromatin states and characterizing their biological function/correlation with large-scale functional datasets. From multiple chromatin markers, Roadmap computed the chromatin states of 127 epigenomes using ChromHMM (v1.10). We downloaded ChromHMM from GREAP (Yang et al. 2022) with 15 states, including TssA (Active TSS), TssAFlnk (Flanking Active TSS), TxFlnk (Transcr. at gene 5” and 3”), etc, more categories can be found in the “Help” page.

Human DNA methylation positions were collected from the ENCODE Human Methylation 450 Bead Chip, which consists of 60 samples with over 500,000 CpG islands. Where beta values greater than 0.6 were considered hypermethylated, while beta values greater than 0.2 and less than 0.6 were considered hypomethylated.

We downloaded eQTLs dataset from VARAdb (Pan et al. 2021) from HaploReg (Ward and Kellis 2012) and PancanQTL (Gong et al. 2018), from GRASP v2.0 (Eicher et al. 2015), GWASdb v2.0 (Li et al. 2016) and the GWAS Catalog (Buniello et al. 2019) for the risk SNPs dataset.

TEs data for mouse and saccharomyces cerevisiae were collected and downloaded from EnhancerAtlas (Gao et al. 2016), and TF ChIP-seq data from GTRD (Yevshin et al. 2017). Mouse SEs data were collected from the dbSUPER (Khan and Zhang 2016).

The final eccDB included human 39,586,081 chromatin accessibility regions, 28,500,150 TFs, 6,370,152 TEs and 393,782 SEs. And 1,515,093 risk SNPs, 10,115,157 eQTLs, 776,369 DNA methylation positions and 37,288,007 chromHMM states were collected. A total of 47,198 TEs on mouse eccDNA, 828 SEs, and 6,294,670 TFs were provided by eccDB. And 739 saccharomyces cerevisiae SEs were also included (Table 1).

### eccDNA score

eccDNAs have an important regulatory function, and to elucidate the strength of its regulatory ability and its importance, we used the scoring algorithm (https://github.com/leishenggit/CircleBase) published by CircleBase (Zhao et al. 2022). Firstly, we downloaded regulatory data (hg19) from CircleBase and converted it to genomic version hg38 using liftOver (Karolchik et al. 2003). The parameter “-F 1” was added to the first step of the scoring algorithm to obtain the number of annotation hits (records) for each eccDNA in each regulatory category. Then run boxcox.py to fit the hits of eccDNA in each regulatory category to a Gaussian model, with the mission median tending to be normally distributed. This was followed by running normalize.py for each category to normalize the scores to [0, 1]. The final score of eccDNA was the average of the normalized scores for each moderation category.

### Chromosomal interaction

Zhu Y *et al*. reported that eccDNAs can function as mobile transcriptional SE elements to promote tumor progression and manifest a potential synthetic aneuploidy mechanism of transcription control in cancer (Zhu et al. 2021). Therefore, eccDNAs chromosomal interactions were divided into intrachromosomal interactions and interchromosomal interactions. If an eccDNA interacts with a chromosome fragment from the same chromosome, that is called an intrachromosomal interaction. If an eccDNA interacts with a chromosomal fragment that is not from the same chromosome, that is called an interchromosomal interaction (Figure 2J). We collected chromosome interaction data (including ChIA-PET, 3C, and Hi-C) from 4DGenome (https://4dgenome.research.chop.edu/; Teng et al. 2015) for three species: Homo sapiens, Mus musculus, and Saccharomyces cerevisiae. The process of human eccDNAs chromosome interactions recognition is as follows: (I) We converted the human interaction regions to human genome hg38 using liftOver (Karolchik et al. 2003). (II) Then selected the human chromosome interaction dataset with “Detection_Method” as “ChIA-PET”, excluding rows with “NA” for “A gene” and “B gene”, “0” for “Confidence_Score1” and “Confidence_Score2”. (III) We converted “Interactor A” region and “Ineractor B” region (Interactor A and Interactor B were two chromosome segments that interact with each other) into nucleic acid sequences by BEDTools getfasta (v2.25.0). (VI) Through BLAST + makeblastdb, the nucleic acid sequences of “Interactor A” and “Interactor B” were constructed as nucleic acid sequence libraries. (V) Each eccDNA was aligned with the “Interactor A” and “Interactor B” nucleic acid libraries by BLAST + blastn. (VI) If the same “Subject_ID (ID of eccDNA similar sequence)” corresponds to more than one “Bit_score”, the “Bit_score” with the high value is retained. (VII) The “Subject_ID” with the top 75% of the “Bit_score” was retained for each eccDNA.

The mouse chromosome interaction dataset was converted to genomic mm10, and the remaining processing steps were identical to those of the human dataset. The saccharomyces cerevisiae chromosome interaction dataset differs from the other two species in that the saccharomyces cerevisiae dataset was selected with a “Detection_Method” of “3C” and “Hi-C” data, and all “Subject_ID” were retained.

Finally, the results of chromosome interactions for all three species were divided into intrachromosomal interactions and interchromosomal interactions. We propose an eccDNA and genome-wide chromosomal interaction prediction method, which is based on sequence similarity and ChIA-PET chromosome interaction data.

For example, if the eccDNA and the region in “Interactor A” are sequence similar (“Subject_ID” is derived from the sequence library built with “Interactor A”), then we hypothesize that there is a potential interaction between the eccDNA and the “Interactor B” corresponding to this “Interactor A”. And if the eccDNA is derived from the same chromosome as “Interactor B”, then it is an intrachromosomal interaction. If the eccDNA is from a different chromosome than “Interactor B”, this is called an interchromosomal interaction.

A total of 3,163,288 human eccDNAs intrachromosomal interactions and 29,454,494 interchromosomal interactions were collected by eccDB. And we collect 79,553 mouse intrachromosomal interactions and 1,417,698 interchromosomal interactions. In addition, we collected a total of 55,579 intrachromosomal interactions and 204,030 interchromosomal interactions in Saccharomyces cerevisiae (Table 1).

### eccDB web interface

The current version of eccDB is developed using MySQL 8.0.21 (http://www.mysql.com) and runs on the Tomcat Web server (http://tomcat.apache.org/). We use JAVA 1.8 (https://www.oracle.com/technetwork/java/index.html) for server-side scripting. The eccDB web interface is designed and built using Bootstrap v3.3.7 (https://v3.bootcss.com), Jquery v2.1.1 (http://jquery.com), Vue (https://cn.vuejs.org/) and Element Plus 2.2.9 (https://element-plus.gitee.io/en-US/). The genomic visualization is accomplished using Integrative Genomics Viewer (IGV) (https://igv.org/). A minimum browser resolution of 1440 × 900 is recommended and use Chrome 105 (or later version), or Firefox 105 (or later version) to achieve the best display.

## Data availability

The public database links used by eccDB are listed in Supplementary Table S2. eccDNA information is available for free download at eccDB (http://www.xiejjlab.bio/eccDB).

## Acknowledgments

This work was supported by grants from the National Natural Science Foundation of China (81871921), the Natural Science Foundation of Guangdong Province-Outstanding Youth Project (2019B151502059), the Basic and Applied Basic Research Programs of Guangdong Province (No. 2018KZDXM033).

## References

Bai X, Shi S, Ai B, Jiang Y, Liu Y, Han X, Xu M, Pan Q, Wang F, Wang Q, et al. 2020. ENdb: a manually curated database of experimentally supported enhancers for human and mouse. Nucleic Acids Res. 48(D1):D51–D57. doi: 10.1093/nar/gkz973

Barretina J, Caponigro G, Stransky N, Venkatesan K, Margolin AA, Kim S, Wilson CJ, Lehár J, Kryukov GV, Sonkin D, et al. 2012. The Cancer Cell Line Encyclopedia enables predictive modelling of anticancer drug sensitivity. Nature. 483(7391):603–7. doi: 10.1038/nature11003

Barrett T, Wilhite SE, Ledoux P, Evangelista C, Kim IF, Tomashevsky M, Marshall KA, Phillippy KH, Sherman PM, Holko M, et al. 2013. NCBI GEO: archive for functional genomics data sets--update. Nucleic Acids Res. 41(Database issue):D991–5. doi: 10.1093/nar/gks1193

Buniello A, MacArthur JAL, Cerezo M, Harris LW, Hayhurst J, Malangone C, McMahon A, Morales J, Mountjoy E, Sollis E, et al. 2019. The NHGRI-EBI GWAS Catalog of published genome-wide association studies, targeted arrays and summary statistics 2019. Nucleic Acids Res. 47(D1):D1005–D1012. doi: 10.1093/nar/gky1120

Camacho C, Coulouris G, Avagyan V, Ma N, Papadopoulos J, Bealer K, Madden TL. 2009. BLAST+: architecture and applications. BMC Bioinformatics. 10:421. doi: 10.1186/1471-2105-10-421

Carithers LJ, Moore HM. 2015. The Genotype-Tissue Expression (GTEx) Project. Biopreserv Biobank. 13(5):307–8. doi: 10.1089/bio.2015.29031.hmm

Chèneby J, Gheorghe M, Artufel M, Mathelier A, Ballester B. 2018. ReMap 2018: an updated atlas of regulatory regions from an integrative analysis of DNA-binding ChIP-seq experiments. Nucleic Acids Res. 46(D1):D267–D275. doi: 10.1093/nar/gkx1092

Corces MR, Granja JM, Shams S, Louie BH, Seoane JA, Zhou W, Silva TC, Groeneveld C, Wong CK, Cho SW, et al. 2018. The chromatin accessibility landscape of primary human cancers. Science. 362(6413):eaav1898. doi: 10.1126/science.aav1898

Deshpande V, Luebeck J, Nguyen ND, Bakhtiari M, Turner KM, Schwab R, Carter H, Mischel PS, Bafna V. 2019. Exploring the landscape of focal amplifications in cancer using AmpliconArchitect. Nat Commun. 10(1):392. doi: 10.1038/s41467-018-08200-y

Dillon LW, Kumar P, Shibata Y, Wang YH, Willcox S, Griffith JD, Pommier Y, Takeda S, Dutta A. 2015. Production of Extrachromosomal MicroDNAs Is Linked to Mismatch Repair Pathways and Transcriptional Activity. Cell Rep. 11(11):1749–59. doi: 10.1016/j.celrep.2015.05.020

Eicher JD, Landowski C, Stackhouse B, Sloan A, Chen W, Jensen N, Lien JP, Leslie R, Johnson AD. 2015. GRASP v2.0: an update on the Genome-Wide Repository of Associations between SNPs and phenotypes. Nucleic Acids Res. 43(Database issue):D799–804. doi: 10.1093/nar/gku1202

ENCODE Project Consortium. 2012. An integrated encyclopedia of DNA elements in the human genome. Nature. 489(7414):57–74. doi: 10.1038/nature11247

Ernst J, Kellis M. 2012. ChromHMM: automating chromatin-state discovery and characterization. Nat Methods. 9(3):215–6. doi: 10.1038/nmeth.1906

Gao T, He B, Liu S, Zhu H, Tan K, Qian J. 2016. EnhancerAtlas: a resource for enhancer annotation and analysis in 105 human cell/tissue types. Bioinformatics. 32(23):3543–3551. doi: 10.1093/bioinformatics/btw495

Gong J, Mei S, Liu C, Xiang Y, Ye Y, Zhang Z, Feng J, Liu R, Diao L, Guo AY, Miao X, Han L. 2018. PancanQTL: systematic identification of cis-eQTLs and trans-eQTLs in 33 cancer types. Nucleic Acids Res. 46(D1):D971–D976. doi: 10.1093/nar/gkx861

Harrow J, Frankish A, Gonzalez JM, Tapanari E, Diekhans M, Kokocinski F, Aken BL, Barrell D, Zadissa A, Searle S, et al. 2012. GENCODE: the reference human genome annotation for The ENCODE Project. Genome Res. 22(9):1760–74. doi: 10.1101/gr.135350.111

Heinz S, Benner C, Spann N, Bertolino E, Lin YC, Laslo P, Cheng JX, Murre C, Singh H, Glass CK. 2010. Simple combinations of lineage-determining transcription factors prime cis-regulatory elements required for macrophage and B cell identities. Mol Cell. 38(4):576–89. doi: 10.1016/j.molcel.2010.05.004

Hnisz D, Abraham BJ, Lee TI, Lau A, Saint-André V, Sigova AA, Hoke HA, Young RA. 2013. Super -enhancers in the control of cell identity and disease. Cell. 155(4):934–47. doi: 10.1016/j.cell.2013.09.053

Hotta Y, Bassel A. 1965. MOLECULAR SIZE AND CIRCULARITY OF DNA IN CELLS OF MAMMALS AND HIGHER PLANTS. Proc Natl Acad Sci U S A. 53(2):356–62. doi: 10.1073/pnas.53.2.356

Hung KL, Yost KE, Xie L, Shi Q, Helmsauer K, Luebeck J, Schöpflin R, Lange JT, Chamorro González R, Weiser NE, et al. 2021. ecDNA hubs drive cooperative intermolecular oncogene expression. Nature. 600(7890):731–736. doi: 10.1038/s41586-021-04116-8

Jiang Y, Qian F, Bai X, Liu Y, Wang Q, Ai B, Han X, Shi S, Zhang J, Li X, et al. 2019. SEdb: a comprehensive human super-enhancer database. Nucleic Acids Res. 47(D1):D235–D243. doi: 10.1093/nar/gky1025

Jung Y, Han D. 2022. BWA-MEME: BWA-MEM emulated with a machine learning approach. Bioinformatics. 2022 Mar 7:btac137. doi: 10.1093/bioinformatics/btac137

Karolchik D, Baertsch R, Diekhans M, Furey TS, Hinrichs A, Lu YT, Roskin KM, Schwartz M, Sugnet CW, Thomas DJ, et al. 2003. The UCSC Genome Browser Database. Nucleic Acids Res. 31(1):51–4. doi: 10.1093/nar/gkg129

Khan A, Zhang X. 2015. dbSUPER: a database of super-enhancers in mouse and human genome. Nucleic Acids Res. 44(D1):D164–71. doi: 10.1093/nar/gkv1002

Kim H, Nguyen NP, Turner K, Wu S, Gujar AD, Luebeck J, Liu J, Deshpande V, Rajkumar U, Namburi S, et al. 2020. Extrachromosomal DNA is associated with oncogene amplification and poor outcome across multiple cancers. Nat Genet. 52(9):891–897. doi: 10.1038/s41588-020-0678-2

Kuleshov MV, Jones MR, Rouillard AD, Fernandez NF, Duan Q, Wang Z, Koplev S, Jenkins SL, Jagodnik KM, Lachmann A, et al. 2016. Enrichr: a comprehensive gene set enrichment analysis web server 2016 update. Nucleic Acids Res. 44(W1):W90–7. doi: 10.1093/nar/gkw377

Kumar P, Dillon LW, Shibata Y, Jazaeri AA, Jones DR, Dutta A. 2017. Normal and Cancerous Tissues Release Extrachromosomal Circular DNA (eccDNA) into the Circulation. Mol Cancer Res. 15(9):1197–1205. doi: 10.1158/1541-7786.MCR-17-0095

Kumar P, Kiran S, Saha S, Su Z, Paulsen T, Chatrath A, Shibata Y, Shibata E, Dutta A. 2020. ATAC-seq identifies thousands of extrachromosomal circular DNA in cancer and cell lines. Sci Adv. 6(20):eaba2489. doi: 10.1126/sciadv.aba2489

Layfield LJ, Willmore C, Tripp S, Jones C, Jensen RL. 2006. Epidermal growth factor receptor gene amplification and protein expression in glioblastoma multiforme: prognostic significance and relationship to other prognostic factors. Appl Immunohistochem Mol Morphol. 14(1):91–6. doi: 10.1097/01.pai.0000159772.73775.2e

Li H, Handsaker B, Wysoker A, Fennell T, Ruan J, Homer N, Marth G, Abecasis G, Durbin R; 1000 Genome Project Data Processing Subgroup. 2009. The Sequence Alignment/Map format and SAMtools. Bioinformatics. 25(16):2078–9. doi: 10.1093/bioinformatics/btp352

Li MJ, Liu Z, Wang P, Wong MP, Nelson MR, Kocher JP, Yeager M, Sham PC, Chanock SJ, Xia Z, et al. 2016. GWASdb v2: an update database for human genetic variants identified by genome-wide association studies. Nucleic Acids Res. 4(D1):D869–76. doi: 10.1093/nar/gkv1317

Liao Z, Jiang W, Ye L, Li T, Yu X, Liu L. 2020. Classification of extrachromosomal circular DNA with a focus on the role of extrachromosomal DNA (ecDNA) in tumor heterogeneity and progression. Biochim Biophys Acta Rev Cancer. 1874(1):188392. doi: 10.1016/j.bbcan.2020.188392

Ling X, Han Y, Meng J, Zhong B, Chen J, Zhang H, Qin J, Pang J, Liu L. 2021. Small extrachromosomal circular DNA (eccDNA): major functions in evolution and cancer. Mol Cancer. 20(1):113. doi: 10.1186/s12943-021-01413-8

Mann L, Seibt KM, Weber B, Heitkam T. 2022. ECCsplorer: a pipeline to detect extrachromosomal circular DNA (eccDNA) from next-generation sequencing data. BMC Bioinformatics. 23(1):40. doi: 10.1186/s12859-021-04545-2

Mei S, Qin Q, Wu Q, Sun H, Zheng R, Zang C, Zhu M, Wu J, Shi X, Taing L, et al. 2017. Cistrome Data Browser: a data portal for ChIP-Seq and chromatin accessibility data in human and mouse. Nucleic Acids Res. 45(D1):D658–D662. doi: 10.1093/nar/gkw983

Møller HD, Pa rsons L, Jørgensen TS, Botstein D, Regenberg B. 2015. Extrachromosomal circular DNA is common in yeast. Proc Natl Acad Sci U S A. 112(24):E3114–22. doi: 10.1073/pnas.1508825112

Møller HD, Ramos-Madrigal J, Prada-Luengo I, Gilbert MTP, Regenberg B. 2020. Near-Random Distribution of Chromosome-Derived Circular DNA in the Condensed Genome of Pigeons and the Larger, More Repeat-Rich Human Genome. Genome Biol Evol. 12(1):3762–3777. doi: 10.1093/gbe/evz281

Møller HD. 2020. Circle -Seq: Isolation and Sequencing of Chromosome-Derived Circular DNA Elements in Cells. Methods Mol Biol. 2119:165–181. doi: 10.1007/978-1-0716-0323-9_15

Noer JB, Hørsdal OK, Xiang X, Luo Y, Regenberg B. 2022. Extrachromosomal circular DNA in cancer: history, current knowledge, and methods. Trends Genet. 38(7):766–781. doi: 10.1016/j.tig.2022.02.007

Pan Q, Liu YJ, Bai XF, Han XL, Jiang Y, Ai B, Shi SS, Wang F, Xu MC, Wang YZ, et al. 2021. VARAdb: a comprehensive variation annotation database for human. Nucleic Acids Res. 49(D1):D1431–D1444. doi: 10.1093/nar/gkaa922

Paulsen T, Kumar P, Koseoglu MM, Dutta A. 2018. Discoveries of Extrachromosomal Circles of DNA in Normal and Tumor Cells. Trends Genet. 34(4):270–278. doi: 10.1016/j.tig.2017.12.010

Peng L, Zhou N, Zhang CY, Li GC, Yuan XQ. 2022. eccDNAdb: a database of extrachromosomal circular DNA profiles in human cancers. Oncogene. 41(19):2696–2705. doi: 10.1038/s41388-022-02286-x

Prada-Luengo I, Krogh A, Maretty L, Regenberg B. 2019. Sensitive detection of circular DNAs at single-nucleotide resolution using guided realignment of partially aligned reads. BMC Bioinformatics. 20(1):663. doi: 10.1186/s12859-019-3160-3

Quinlan AR, Hall IM. 2010. BEDTools: a flexible suite of utilities for comparing genomic features. Bioinformatics. 26(6):841–2. doi: 10.1093/bioinformatics/btq033

Shibata Y, Kumar P, Layer R, Willcox S, Gagan JR, Griffith JD, Dutta A. 2012. Extrachromosomal microDNAs and chromosomal microdeletions in normal tissues. Science. 336(6077):82–6. doi: 10.1126/science.1213307

Tang Z, Kang B, Li C, Chen T, Zhang Z. 2019. GEPIA2: an enhanced web server for large-scale expression profiling and interactive analysis. Nucleic Acids Res. 47(W1):W556–W560. doi: 10.1093/nar/gkz430

Teng L, He B, Wang J, Tan K. 2015. 4DGenome: a comprehensive database of chromatin interactions. Bioinformatics. 31(15):2560–4. doi: 10.1093/bioinformatics/btv158

Wang F, Bai X, Wang Y, Jiang Y, Ai B, Zhang Y, Liu Y, Xu M, Wang Q, Han X, et al. 2021. ATACdb: a comprehensive human chromatin accessibility database. Nucleic Acids Res. 49(D1):D55–D64. doi: 10.1093/nar/gkaa943

Wang K, Tian H, Wang L, Wang L, Tan Y, Zhang Z, Sun K, Yin M, Wei Q, Guo B, et al. 2021. Deciphering extrachromosomal circular DNA in Arabidopsis. Comput Struct Biotechnol J. 19:1176–1183. doi: 10.1016/j.csbj.2021.01.043

Wang M, Chen X, Yu F, Ding H, Zhang Y, Wang K. 2021. Extrachromosomal Circular DNAs: Origin, formation and emerging function in Cancer. Int J Biol Sci. 17(4):1010–1025. doi: 10.7150/ijbs.54614

Wang S, Tian F, Qiu Y, Liu X. 2010. Bilateral similarity function: a novel and universal method for similarity analysis of biological sequences. J Theor Biol. 265(2):194–201. doi: 10.1016/j.jtbi.2010.04.013

Wang T, Zhang H, Zhou Y, Shi J. 2021. Extrachromosomal circular DNA: a new potential role in cancer progression. J Transl Med. 19(1):257. doi: 10.1186/s12967-021-02927-x

Ward LD, Kellis M. 2012. HaploReg: a resource for exploring chromatin states, conservation, and regulatory motif alterations within sets of genetically linked variants. Nucleic Acids Res. 40(Database issue):D930–4. doi: 10.1093/nar/gkr917

Wu S, Turner KM, Nguyen N, Raviram R, Erb M, Santini J, Luebeck J, Rajkumar U, Diao Y, Li B, et al. 2019. Circular ecDNA promotes accessible chromatin and high oncogene expression. Nature. 575(7784):699–703. doi: 10.1038/s41586-019-1763-5

Xu G, Li JY. 2018. CDK4, CDK6, cyclin D1, p16(INK4a) and EGFR expression in glioblastoma with a primitive neuronal component. J Neurooncol. 136(3):445–452. doi: 10.1007/s11060-017-2674-7

Yevshin I, Sharipov R, Valeev T, Kel A, Kolpakov F. 2017. GTRD: a database of transcription factor binding sites identified by ChIP-seq experiments. Nucleic Acids Res. 45(D1):D61–D67. doi: 10.1093/nar/gkw951

Yang Y, Qian F, Li X, Li Y, Zhou L, Wang Q, Zhou X, Zhang J, Song C, Yu Z, et al. 2022. GREAP: a comprehensive enrichment analysis software for human genomic regions. Brief Bioinform. 2022 Aug 12:bbac329. doi: 10.1093/bib/bbac329

Zhang P, Peng H, Llauro C, Bucher E, Mirouze M. 2021. ecc_finder: A Robust and Accurate Tool for Detecting Extrachromosomal Circular DNA From Sequencing Data. Front Plant Sci. 12:743742. doi: 10.3389/fpls.2021.743742

Zhao X, Shi L, Ruan S, Bi W, Chen Y, Chen L, Liu Y, Li M, Qiao J, Mao F. 2022. CircleBase: an integrated resource and analysis platform for human eccDNAs. Nucleic Acids Res. 50(D1):D72–D82. doi: 10.1093/nar/gkab1104

Zhu Y, Gujar AD, Wong CH, Tjong H, Ngan CY, Gong L, Chen YA, Kim H, Liu J, Li M, et al. 2021. Oncogenic extrachromosomal DNA functions as mobile enhancers to globally amplify chromosomal transcription. Cancer Cell. 39(5):694-707.e7. doi: 10.1016/j.ccell.2021.03.006

Zuo S, Yi Y, Wang C, Li X, Zhou M, Peng Q, Zhou J, Yang Y, He Q. 2022. Extrachromosomal Circular DNA (eccDNA): From Chaos to Function. Front Cell Dev Biol. 9:792555. doi: 10.3389/fcell.2021.792555

